# Disruption of the mitochondrial gene *orf352* partially restores pollen development in cytoplasmic male sterile rice

**DOI:** 10.1101/2021.02.24.432723

**Authors:** Shiho Omukai, Shin-ich Arimura, Kinya Toriyama, Tomohiko Kazama

## Abstract

Plant mitochondrial genomes sometimes carry cytoplasmic male sterility (CMS)-associated genes. These genes have been harnessed in agriculture to produce high-yielding F_1_ hybrid seeds in various crops. The gene *orf352* was reported to be an RT102-type CMS gene in rice (*Oryza sativa*), although a causal demonstration of its role in CMS is lacking. Here, we employed mitochondrion-targeted transcription activator-like effector nucleases (mitoTALENs), to knock out *orf352* from the mitochondrial genome in the cytoplasmic male sterile rice RT102A. We isolated 18 independent transformation events in RT102A that resulted in genome editing of *orf352*, including its complete removal from the mitochondrial genome in several plants. Sequence analysis around the mitoTALEN target sites revealed the repair of their induced double-strand breaks via homologous recombination. Near the 5ʹ target site, repair involved sequences identical to *orf284*, while repair of the 3ʹ target site yielded various new sequences that generated new chimeric genes consisting *orf352* fragments. Plants with a new mitochondrial gene encoding amino acids 179 to 352 of ORF352 exhibited the same shrunken pollen grain phenotype as RT102A, whereas plants either lacking *orf352* or harboring a new gene encoding amino acids 211 to 352 of ORF352 showed partial rescue of pollen viability and germination, although they failed to set seed. These results demonstrated that disruption of *orf352* partially restored pollen development, indicating that amino acids 179 to 210 from ORF352 may contribute to the establishment of pollen abortion.

## Introduction

Cytoplasmic male sterility (CMS) was first reported in maize (*Zea mays*) (Rhoades, 1931). In rice (*Oryza sativa*), the first report of a cytoplasmic effect on male function was reduction of seed fertility in the progeny of the first backcross between *O. rufipogon* and *O. sativa* (Katsuo and Mizushima, 1958). In agriculture, CMS is harnessed to produce high-yielding F_1_ hybrid seeds in various crops. Several genes in the mitochondrial genome cause dysfunction of pollen development, resulting in male sterility. These genes are ordinarily named *orf* (*open reading frame*) followed by a unique number, indicating the number of amino acids in encoded proteins, and they are associated with cytoplasmic male sterility (CMS). CMS-causative genes in the mitochondrial genome are broadly referred to as CMS-associated genes because a direct causal relationship between *orf* and CMS is, for the most part, technically out of reach at present. CMS-associated genes have been reported in many plants (Hanson & Bentolila, 2004). Examples of such genes in rice include *orf79* in BT-type CMS (Iwabuchi et al., 1993; Akagi et al., 1994), *orfH79* in HL-type CMS (Yi P, 2002), *orf352* in WA-type CMS (Bentolila & Stefanov, 2012; Luo et al., 2013), and CW-*orf307* in CW-type CMS (Fujii et al., 2010). Similarly, in Brassicaceae, cybrid between rapeseed (*Brassica napus*) and radish (*Raphanus sativus*) led to the identification of *orf125* in Kosena CMS (Iwabuchi et al., 1999), and *orf138* in Ogura CMS (Grelon et al., 1994).

We recently employed transcription activator-like effector nucleases (TALENs) to delete the mitochondrial CMS-associated gene *orf79* in BT-type CMS rice (Kazama et al., 2019). TALENs, which make genome editing possible, usually comprise the left TALEN and right TALEN, each containing a designer DNA-binding domain and a nuclease domain. TALENs typically function as obligate heterodimers, and are often used to induce double-strand breaks (DSBs) at or near the region recognized by the DNA-binding domain. These are then repaired by imperfect nonhomologous end-joining (NHEJ) repair in the nuclear genome, introducing small deletions or insertions in the process. We designed a mitoTALEN vector to add a mitochondrial targeting signal to a TALEN. The mitoTALEN construct is integrated into the nuclear genome via Agrobacterium (*Agrobacterium tumefaciens*)-mediated transformation, thereby allowing the mitochondrial genome to be edited without the need for direct transformation of the organellar genome. Using this method, we successfully deleted *orf79* in BT-type CMS rice and restored fertility, illustrating the power of mitoTALENs as tools to reveal the role of a CMS-associated gene.

Traditionally, the presence of a CMS-causative gene is revealed by outcrossing a line to separate the mitochondrial CMS locus and the nuclear *restorer of fertility* (*Rf*) gene that normally suppresses the effects of the CMS-causative gene. In this study, we used RT102-type CMS rice, which is derived from the cytoplasm of the wild rice *O. rufipogon* Griff., accession W1125 (Motomura et al., 2003). The CMS line RT102A carries the cytoplasm of RT102 and the nucleus of the fertile *japonica* cultivar, Taichung 65 (T65). Pollen grains in RT102A are sterile, with two different morphologies: approximately 83% are shrunken and cannot be stained by Lugol’s iodine, while the remaining 17% are spherical and stained strongly (Okazaki et al., 2013). The shrunken pollen phenotype seen in RT102A is like that of WA-CMS, in which microspores abort at the early uninucleate microspore stage, right after meiosis (Li, 2007). We previously sequenced and assembled the RT102-type mitochondrial genome into a single circular molecule of 502,250 bp and identified the new chimeric gene, *orf*352, as a CMS-associated gene (Okazaki et al., 2013). The protein encoded by *orf*352 was almost identical to WA352, a CMS-associated protein in WA-type CMS (Bentolila & Stefanov, 2012; Luo et al., 2013; Tang et al., 2017), with only five nucleotide differences resulting in four amino acid substitutions. The *orf352* gene appeared to comprise a fragment identical to a part of *orf284*, and a fragment with similarity to *orf288*, both of which encode hypothetical proteins. Previously, *orf284* and *orf288* have been reported to be involved in the generation of CMS-associated genes (Tang et al., 2017). Functional evidence for the role of *orf352*/*WA352* in RT102-/WA-type CMS as a CMS-causative gene has not been described.

Here, we used loss-of function analysis to assess the contribution of the mitochondrial chimeric gene *orf352* to CMS in the CMS line RT102A. We generated two mitoTALEN vectors to target *orf352* and introduced them into the CMS line RT102A. We describe the characterization of the genotypes and phenotypes and phenotypes associated with genome editing of *orf352* via mitoTALENs and suggest that disruption of *orf352* partially restores pollen development.

## Results

### Nuclear transformation of RT102-type CMS rice with TALENs targeting mitochondrial *orf352*

Mitochondrial *orf352* was previously reported to be a CMS-associated gene in the RT102-type CMS rice, RT102A (Okazaki et al., 2013). To determine what contribution, if any, *orf352* might make to the male sterile phenotype displayed in RT102A, we searched for a sequence unique to *orf352* before designing TALENs. The *orf*352 coding region comprises 1,059 nucleotides (nt) and is divided into three fragments based on their homology to other mitochondrial genes. The first *orf352* fragment is 297 nt in length and is identical to *orf284*, while the last 562 nt of *orf352* (from 498–1,059 nt, where the first adenine from the translation start codon is counted as 1) has 97% sequence identity with *orf288* (Fig. 1A, B). The middle *orf352* fragment (298–497 nt) is specific to the *orf352*. We therefore selected this region as a target sequence for mitoTALENs (Fig. S1A). We generated two mitoTALEN vectors, pTAL1 and pTAL2, harboring the left and the right part of each TALEN in each mitoTALEN vector. Each mitoTALEN vector was introduced into RT102A rice calli via Agrobacterium (*Agrobacterium tumefaciens*)-mediated transformation. We confirmed stable insertion of the transgene into the plant genome by PCR amplification of an introduced hygromycin resistance gene (*HPT*) present on the T-DNA (Fig. S1B). Six primary (T_0_) transgenic plants harboring pTAL1, and 12 harboring pTAL2, were generated.

**Figure 1.**
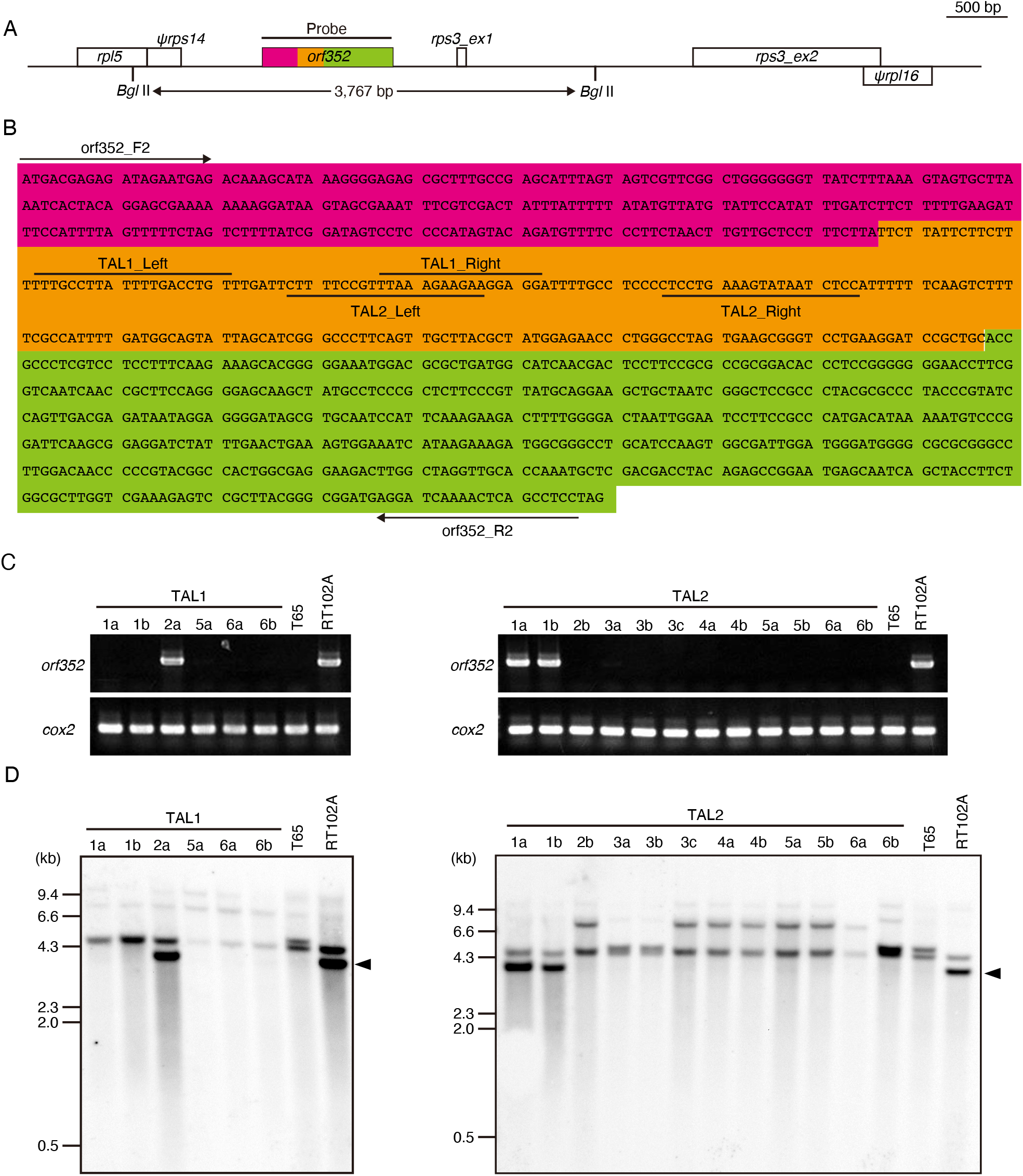
Introduction of the mitoTALENs TAL1 and TAL2, targeting *orf352* in the RT102-type CMS rice (RT102A). **(A)** Schematic illustration of the target gene *orf352* and the neighboring region in the RT102-type mitochondrial genome. Pink, region identical to *orf284*; green, region homologous to *orf288*; *orf352*-specific sequences are shown in orange. **(B)** Nucleotides sequences of the *orf352* coding region. Background colors are as in (A). Primers for PCR analysis (*orf352*_F2, *orf352*_R2) are shown, as well as each TAL binding site. **(C)** Genotyping of transgenic plants by PCR over the *orf352* region. T65 is a fertile *japonica* rice cultivar that lacks *orf352* and served as a negative control. **(D)** Southern blot analysis of transgenic plants with TAL1 and TAL2-specific probes. Arrowheads indicate signals corresponding to a 3.8-kb *orf352* fragment.

The introduction of a mitoTALEN targeting *orf79* has been reported to be accompanied by the disappearance of *orf79* from the mitochondrial genome of transformants (Kazama et al., 2019). To determine whether *orf352* might be similarly deleted by mitoTALENs, we tested genomic DNA from transformants for the presence of *orf352* by PCR and Southern blot analysis (Fig. 1C, D). The PCR primers *orf352*_F2 and *orf352*_R2 should amplify the full-length *orf352* coding region from RT102A genomic DNA, as well as in any transformants retaining the entire *orf352* sequence. However, with the exception of TAL1-2a, we failed to amplify *orf352* from most plants transformed with pTAL1. Likewise, except for plants TAL2-1a and TAL2-1b, most plants carrying the T-DNA derived from pTAL2 were negative for *orf352*, as tested by PCR amplification (Fig. 1C). We amplified full-length *orf352* from RT102A genomic DNA; since RT102A and the transformants TAL1-2a, TAL2-1a, and TAL2-1b plants shared the same band pattern for the PCR products after electrophoresis on agarose gel, we hypothesized that the *orf352* PCR-positive plants might not carry any alteration at *orf352*. To investigate this possibility, we sequenced the PCR products obtained above and compared their sequences with the RT102A mitochondrial genome. The sequences were identical, confirming the absence of rearrangements around the mitoTALEN target sites. We also verified the presence or absence of the *orf352* by Southern blot analysis using total genomic DNA extracted from leaf blades, and a probe spanning the *orf352* region, as indicated in Figure 1A. We detected a 3.8-kb band in RT102A, as expected for the intact *orf352* genomic fragment. However, we did not detect a band of the same size or intensity in the fertile *japonica* cultivar, Taichung 65 (T65), which does not carry *orf352* in its mitochondrial genome. Five of the TAL1 transgenic plants (TAL1-1a, 1b, 5a, 6a, and 6b) and 10 of the TAL2 transgenic plants (TAL2-2b, 3a, 3b, 3c, 4a, 4b, 5a, 5b, 6b, and 6c) similarly showed no signal specific for *orf352* (Fig. 1D). The same transgenic plants were also negative for *orf352* by PCR (Fig. 1C). However, the three *orf352*-positive transgenic plants by PCR (TAL1-2a, TAL2-1a, and TAL2-1b) had a strong hybridization signal at the same size as that of RT102A. These results indicate that *orf352* is intact in the mitochondrial genome in the transgenic plants.

### Aberrant homologous recombination events in mitochondrial genomes edited at *orf352*

Previous analysis of mitoTALEN-mediated genome editing of the mitochondrial gene *orf79* revealed that the double-strand break (DSB) introduced by mitoTALENs is repaired by two independent homologous recombination events at recombination sites A and B on either side of the target site (Kazama et al., 2019) (Fig. S2). Indeed, the DSB is repaired via recombination between site A (or B) and site Aʹ (or Bʹ), which refers to a region anywhere in the mitochondrial genome with high sequence similarity to site A (or B) (Fig. S2). During DSB repair, the region between recombination sites A and B, including the target site itself, is deleted from the mitochondrial genome. The homologous recombination occurs non-reciprocally and is accompanied by replication, so finally, DNA molecules containing the site A/Aʹ and B/Bʹ will be duplicated (Fig. S2).

To determine the extent of the deletion around *orf352* extended, we designed specific primer pairs to PCR-amplify eight fragments mapping up to 1.4-kb upstream and up to 3.8-kb downstream of *orf352* (Fig. S3A). We successfully amplified the four amplicons upstream of *orf352* (regions 1–4), as well as the two downstream amplicons (regions 7 and 8), in all transgenic plants, with the exception of transgenic plants TAL1-1a, TAL1-1b, and TAL1-5a. This indicated that these three transformants carry larger deletions around *orf352* than any of the other transgenic plants (Fig. S3B). To rule out the possibility that DSBs have been repaired by end-joining rather than homologous recombination, we attempted to amplify across the entire region around *orf352* (labeled as region 9 in Fig. S3A) with the forward primer from region 3 and the reverse primer from region 7. End-joining repair would be expected to cause a deletion or an insertion, which would be apparent on an agarose gel compared with the product from RT102A. However, we failed to obtain a PCR amplicon from *orf352*-editied plants, even though all *orf352*-positive plants exhibited an amplicon of the same size as RT102A (Fig. S3B). This result indicates that the DSBs were not repaired by end-joining.

We have noticed previously that mitoTALEN-induced DSBs are repaired via homologous recombination at an arbitrary recombination site at an arbitrary position from the mitoTALEN target site (Kazama et al., 2019). To better understand the recombination events in each transformant, we turned to fusion primer and nested integrated PCR (FPNI-PCR), which is a modified method of thermal asymmetric interlaced (TAIL)-PCR (Wang et al., 2011). We then sequenced the FPNI-PCR amplicons using primers flanking the recombination sites and compared the resulting nucleotide sequences to the RT102-type mitochondrial genome using BLAST to identify the template used during repair (Fig. 2). Our query sequences sometimes showed high sequence similarity to two regions in the single master circle of the RT102-type mitochondrial genome, likely because of the large duplication it carries. We identified six distinct types of recombination on the 3ʹ mitoTALEN-target site (Fig. 2A and Fig. S4). In Type 1 (TAL1-1a), Type 2 (TAL1-1b), and Type 3 (TAL1-5a) recombination events, the entire *orf352* coding region was lost. Based on BLAST results, the DSB from the Type 1 recombination event was repaired via homologous recombination between a 17-bp region downstream of *orf352* (267450 nt to 267,466 nt in the RT102-type mitochondrial genome, GenBank AP012528), and a distant 17-bp region (221,704–221,719 nt) (Fig. 2B). In the Type 2 recombination event (TAL1-1b), the DSB was repaired between a 17-bp region (267,510–267,526 nt) downstream of *orf352*, and one of two possible distant 17-bp regions (110,198–110,214 nt and 441,826–441,842 nt). Homologous recombination in Type 3 plants involved a 100-bp region downstream of *orf352* (267,776–267,845 nt) and a distant 96-bp region (209,174–209,269 nt) as template for repair. In plants with Type 4 (TAL1-6a and Type1-6b), Type 5 (TAL2-2b, TAL2-3c, TAL2-4a, TAL2-4b, TAL2-5a, TAL2-5b, and TAL2-6b), and Type 6 (TAL2-3a, TAL2-3b, and TAL2-6c) repair, the recombination sites were located within the *orf352* (268,339–268,354 nt) and a distant 16-bp region (206,014–206,029 nt). The recombination site for Type 5 and Type 6 repair is identical, but Type 5 repair appears to include an additional recombination event 95-bp upstream of the common recombination site for Type 5 and Type 6 repair (Fig. 4S).

**Figure 2.**
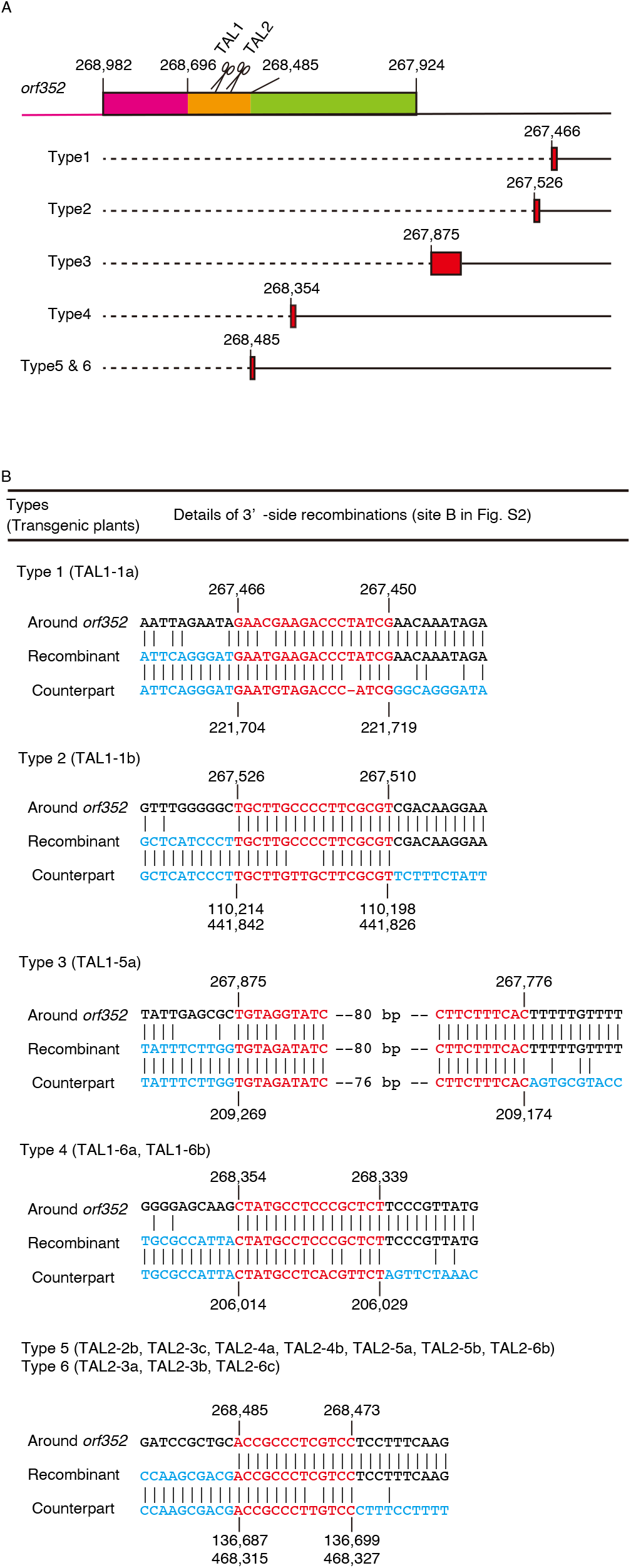
Double-strand breaks induced by mitoTALENs are repaired via homologous recombination. **(A)** Schematic diagram of the various recombination types observed at site B (the principle behind recombination is illustrated in Fig. S2). Top, genetic structure around the *orf352* coding region. Pink, region identical to *orf284*; green, region homologous to *orf288*; orange, *orf352*-specific sequences. Scissors indicate mitoTALENs (TAL1 [left] and TAL2 [right]). Bottom, positions (in bp) and structure of the recombination event at site B in each type of transformant. Red filled boxes indicate the recombination sites B. **(B)** Sequence comparisons between the RT102A mitochondrial genome and each recombinant type.

Although repair on the 3ʹ side of the mitoTALEN target site varied across all transgenic plants, recombination at the 5ʹ site (recombination site A, as illustrated in Fig. S2) made use of the same site in all *orf352*-edited transgenic plants (Fig. 3 and Fig. S5). Results from BLAST searched against the RT102-type mitochondrial genome indicated that repair took place between the first half of *orf352* (268,696–269,254 nt, with homology to *orf284* and its promoter region) and the region 5ʹ of *orf284* in the mitochondrial genome (115,367–115,925 nt, and 446,995–447,553 nt) (Fig. 3 and Fig. S5). The identical sequence of 559 bp are illustrated in Fig. S2 as the recombination sites A and Aʹ. Repair via homologous recombination at this site resulted in the duplication of *orf284*, with each *orf284* having a distinct upstream sequence, which we confirmed by PCR in all *orf352*-edited plants. We also independently confirmed the duplication by Southern blot analysis, which showed two bands when using *orf284* (amplified with P4 and P5) as a probe (Fig. 3). Like the *orf79* in BT-type CMS, DSBs in *orf352* induced by mitoTALENs were repaired via homologous recombination and we successfully obtained *orf352*-edited plants.

**Figure 3.**
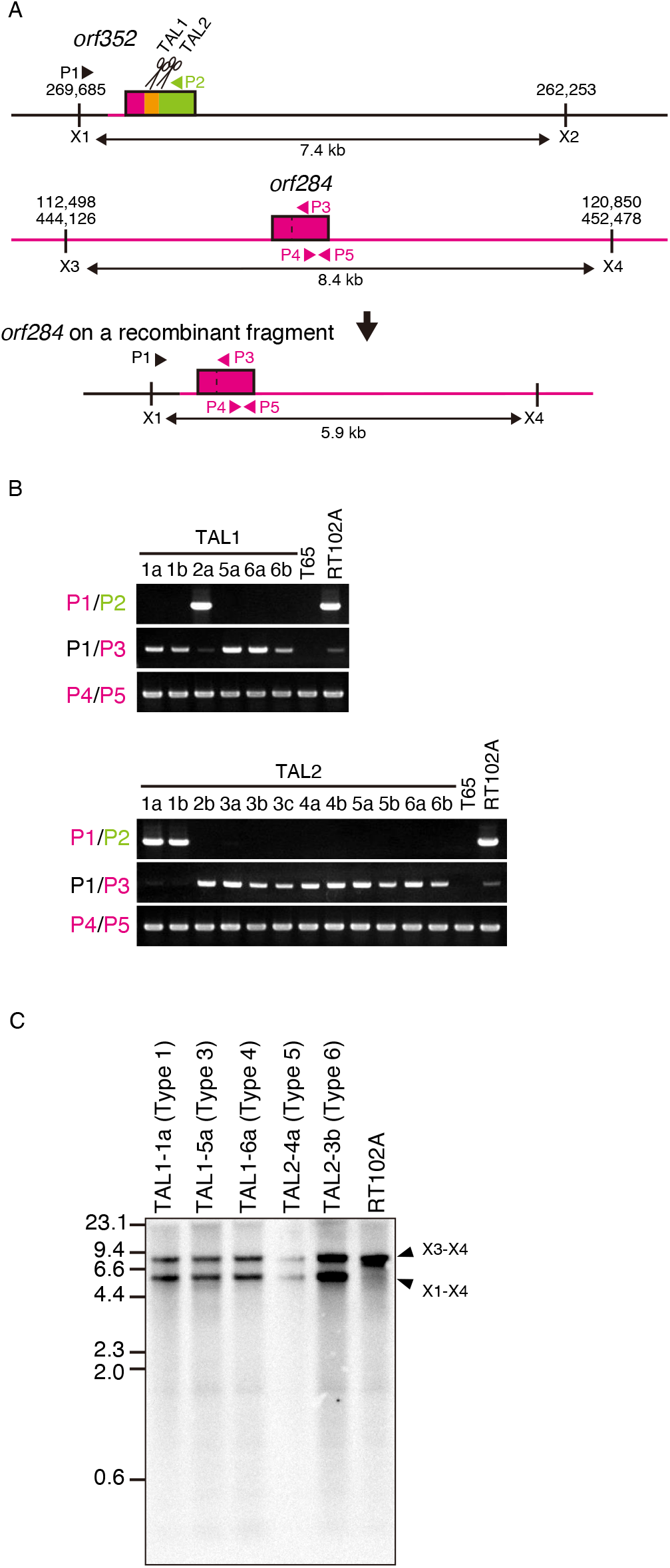
Double-strand breaks induced by mitoTALENs are repaired by identical homologous recombination in all *orf352*-edited plants. **(A)** Schematic diagram of double-strand break repair at the 5ʹ side of the target site. Top, genomic structure around the *orf352* open reading frame. Pink, region identical to *orf284*; green, region homologous to *orf288*; orange, *orf352*-specific sequences. Scissors indicate mitoTALENs (TAL1 and TAL2). Middle, genomic structure around *orf284*. Bottom, genomic structure of new recombinants. Black line, region originating from around *orf352*. P1–P5, primer positions for genotyping PCR; X1–X4, positions of *Xho* I restriction sites. **(B)** PCR analysis of new recombinants. T65 is a fertile *japonica* rice that lacks *orf352*, which serves as a negative control. RT102A serves as a positive control for the presence of *orf352*. **(C)** Duplication of *orf284* during repair via homologous recombination, as detected by Southern blot analysis hybridizing on genomic DNA digested with *Xho* I. The probe was synthesized using primers P4 and P5.

### Partial restoration of pollen development, but not seed set, via genome editing of *orf352*

Genome editing of *orf79* has been shown to restore fertility of BT-type CMS rice (Kazama et al., 2019). To test the effects of *orf352* editing on fertility, we calculated the seed setting rate for all transgenic plants (Table 1). The seed setting rate for the fertile cultivar T65 was close to 90%, whereas the CMS line RT102A was completely sterile, as expected (Okazaki et al., 2013). Contrary to our expectations, all *orf352*-edited plants were also completely sterile (Table 1), indicating that *orf352* is unlikely to be the cause of RT102-type CMS. We expanded our analysis of the consequences of *orf352* genome editing to pollen viability and pollen tube elongation. Most pollen grains from cultivar T65 were stained by Lugol’s iodide solution, demonstrating their viability, whereas most RT102A pollen grains were shrunken and few were stained by Lugol’s (Okazaki et al., 2013) (Fig. 4). The few spherical pollen grains in RT102A appeared normal in appearance but did not germinate when placed on stigmas. By contrast, pollen grains from cultivar T65 germinated on stigmas and their tubes elongated, a clear sign of viable and fertile pollen (Fig. 4).

**Table 1.** Seed setting rate for *orf352*-editied plants

| Line | No. of spikelets/panicles | No. of filled<br>grains | Seed setting<br>rate (%) | *orf352*<sup>1</sup> |
| --- | --- | --- | --- | --- |
| T65 | 355/6 | 308 | 87.0 ± 2.9 | — |
| RT102A | 154/3 | 0 | 0.0 ± 0.0 | + |
| TAL1-1a (Type 1) | 226/10 | 0 | 0.0 ± 0.0 | — |
| TAL1-1b (Type 2) | 154/3 | 0 | 0.0 ± 0.0 | — |
| TAL1-5a (Type 3) | 291/3 | 0 | 0.0 ± 0.0 | — |
| TAL1-6a (Type 4) | 542/10 | 0 | 0.0 ± 0.0 | Disrupted |
| TAL2-4a (Type 5) | 112/2 | 0 | 0.0 ± 0.0 | Disrupted |
| TAL2-3b (Type 6) | 330/8 | 0 | 0.0 ± 0.0 | Disrupted |

<sup>1</sup> —, *orf352* ORF is absent in the mitochondrial genome; +, *orf352* ORF is present in the mitochondrial genome

**Figure 4.**
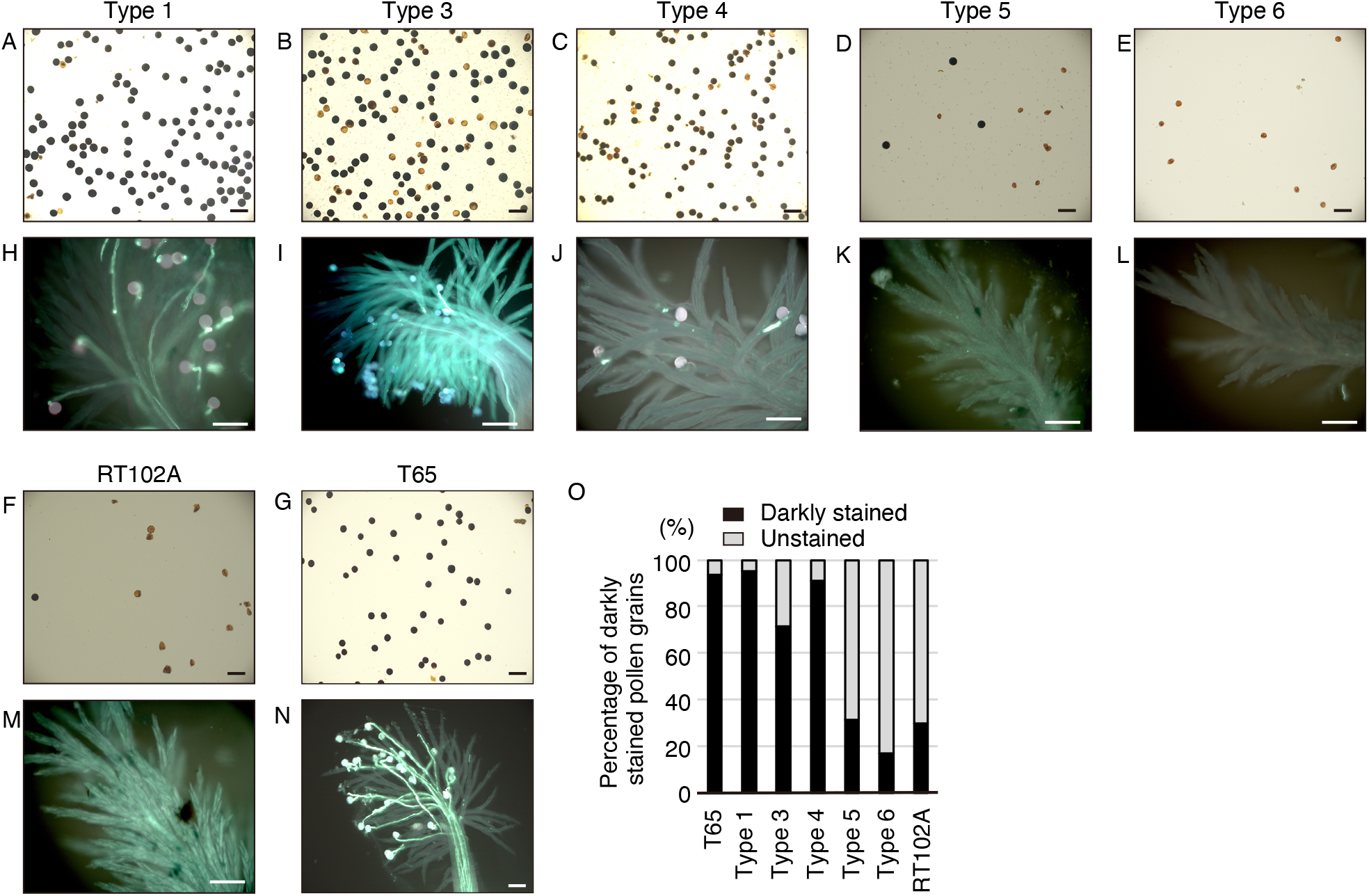
Disruption of *orf352* partially restores pollen development but does not restore pollen fertility. **(A–G)** Pollen grains viability in the various recombinant types, as determined by staining with Lugol’s iodine (1% I_2_-KI). **(H–N)** Pollen germination and pollen tube elongation on stigmas, observed by staining pollen tubes with aniline blue. **(O)** Percentage of darkly stained pollen grains (viable) in each recombinant type. Scale bars = 200 μm (A–G) and 100 μm (H–N).

Having established that the control cultivars displayed their expected fertility phenotypes, we next checked the pollen phenotypes for all but one *orf352*-edited plants (one Type 2 plant became blighted before anthesis). In Type 1 (TAL1-1a), Type 3 (TAL1-5a), and Type 4 (TAL1-6a) plants, the percentage of darkly stained pollen grains was similar to that seen in the fertile cultivar T65, with values of 96%, 84%, and 91%, respectively (Fig. 4). By contrast, Type 5 and Type 6 plants showed a severe reduction in pollen viability, with a percentage of dark pollen grains of 31% and 17%, respectively – lower than or similar to the 30% viable pollen seen in RT102A (Fig. 4). In all transgenic plants, far fewer pollen grains germinated on stigmas relative to pollen grains from T65. Although pollen tubes did not start to elongate, they quickly stopped progressing along the stigma, either just after germination or after reaching about halfway down the stigma. Almost all pollen grains from Type 5 and Type 6 plants were empty and shrunken, similar to RT012A pollen, and none germinated, as predicted. To qualify the pollen phenotypes of *orf352*-edited plants, we counted the number of darkly stained pollen grains. Over 60% of pollen grains from Type 1, Type 3, and Type 4 plants were stained, indicative of starch reserves. In these transgenic plants, pollen viability therefore partially recovered relative to that observed in Type 5 or Type 6 plants, or in the CMS line RT102A. Type 5 and Type 6 plants, however, showed no restoration of pollen viability. These data indicate that *orf352* is involved in inhibition of pollen development, and other unidentified gene(s) are involved in inducing male sterility in RT102A.

### Creation of new predicted genes near recombination sites

Targeting *orf352* with mitoTALENs induced the formation of DSBs, which were repaired via homologous recombination between homologous sequences near the target and anywhere in the mitochondrial genome (Fig. 2, Fig. 3, and Fig. S2). New genes may thus emerge at the recombination sites. To explore this possibility, we PCR-amplified the recombined DNA fragments containing either recombination site A/Aʹ or B/Bʹ (Fig. S2) and determined their sequences. A new gene was defined as the longest gene not previously present in the mitochondrial genome. At the 5ʹ recombination site (indicated as A/Aʹ in Fig. S2), all *orf352*-edited plants shared the same duplicated gene, *orf284*. We predicted several new genes at the 3ʹ recombination sites, which we previously showed to be associated with homologous recombination events (Fig. S4 and Table 2). One new gene was observed in each of Type 1 and Type 2 recombination repair events, encoding of 77 and 47 amino acids, respectively, which we named *orf77* (Type 1) and *orf47* (Type 2). Both are completely new *open reading frame*s with no similarity to other genes or known genes; however, the mitochondrial genome of Type 3 plants harbored a new gene encoding a protein of 108 amino acids, which included the entire ATP synthase subunit 9 proteins (ATP9). Recombination in Type 4 plants created an *open reading frame* encoding a 142-aa protein, comprising two new amino acids added to the N terminus of a fragment of ORF352 (from 215–352 aa). We named this gene as *orf142*. Recombination events in Type 5 and Type 6 plants created the same gene, *orf174*, encoding a protein comprising 179–352 aa of ORF352. Additional recombination upstream of *orf174* occurred in Type 5, so Type 5 and Type 6 are distinguished from each other (Fig. S4). While pollen viability was not restored in Type 5 or Type 6 plants, it was in other transgenic plants, raising the possibility that a fragment comprising 179–210 aa of ORF352 is responsible for pollen development inhibition.

**Table 2.**
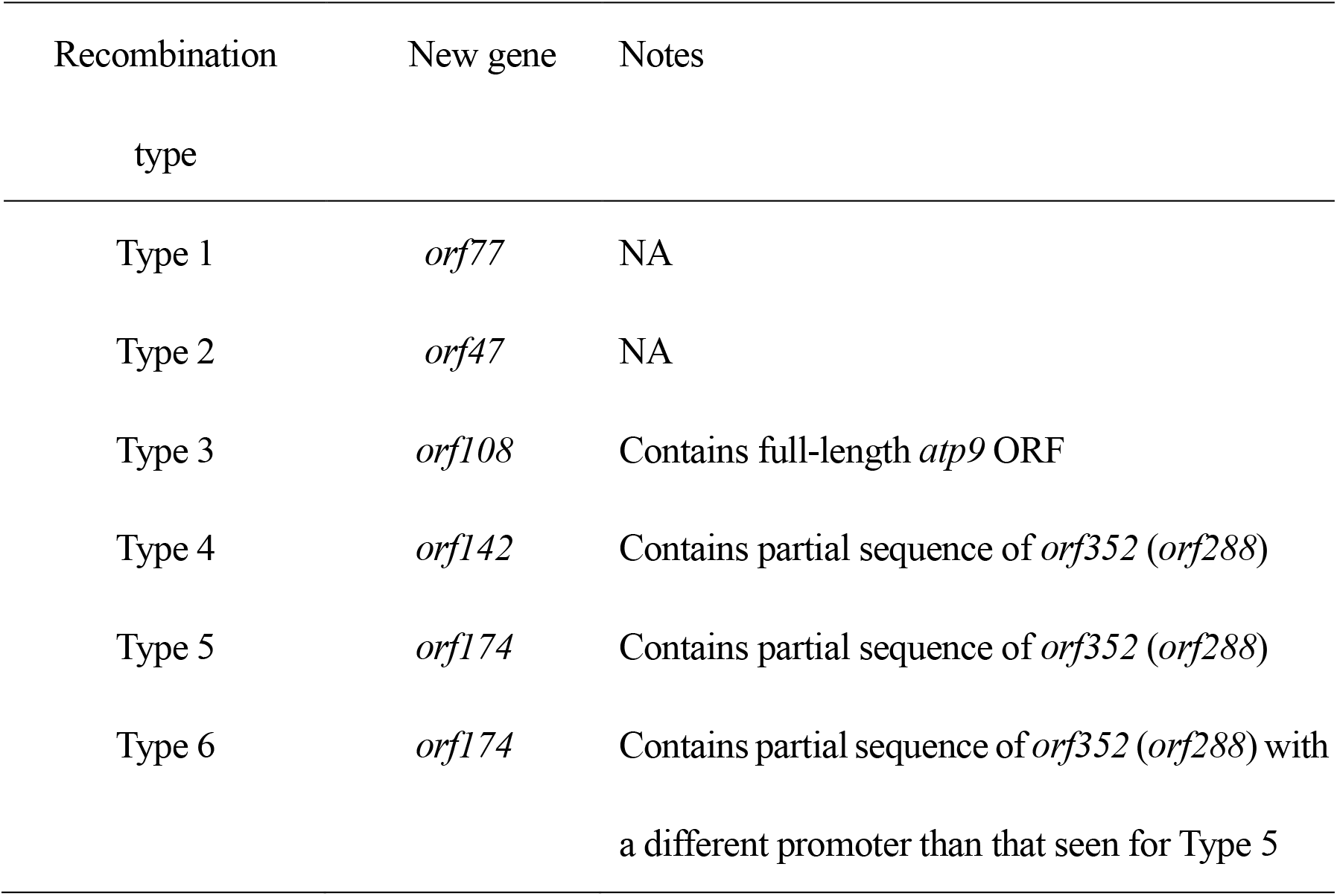
New loci resulting from recombination at the 3ʹ site.

| Recombination<br>type | New gene | Notes |
| --- | --- | --- |
| Type 1 | *orf77* | NA |
| Type 2 | *orf47* | NA |
| Type 3 | *orf108* | Contains full-length *atp9* ORF |
| Type 4 | *orf142* | Contains partial sequence of *orf352* (*orf288*) |
| Type 5 | *orf174* | Contains partial sequence of *orf352* (*orf288*) |
| Type 6 | *orf174* | Contains partial sequence of *orf352* (*orf288*) with<br>a different promoter than that seen for Type 5 |

## Discussion

Hybrid vigor confers improved performance and increased yield to F_1_ hybrid plants over either parent. Global seed markets for most vegetables and major crops such as maize (*Zea mays*), sorghum (*Sorghum bicolor*), and rice, are thus dominated by F_1_ hybrid seeds (Hochholdinger & Baldauf, 2018). To avoid self-pollination that would lead to the production of non-hybrid seed, male-sterile plants are routinely employed as the female parent in crosses; thus, CMS has considerable agronomic importance. In rice, wild abortive (WA)-type CMS is mainly used to produce hybrid seed of *indica* varieties (Huang et al., 2014), and is associated with shrunken pollen grains at the flowering stage that cannot be stained with Lugol’s iodine solution. The fertility of this CMS can be restored by the nucleus-encoded fertility restorer genes, *Rf3* and *Rf4*. Sequencing the mitochondrial genome in WA-type CMS plants identified *orf352* as a putative CMS-causative gene, named *WA352*. The WA352 protein has been reported to accumulate preferentially in the anther tapetum and interacts with the COX11 subunit of a respiration complex IV, resulting in ROS production and programmed cell death, which eventually leads to microspore abortion at the early microspore stage (Luo et al., 2013). The restorer of fertility protein RF4 is thought to function at the post-transcriptional level, while RF3 acts post-translationally to suppress the accumulation of WA352. The cloning of *Rf4* revealed that RF4 is a pentatricopeptide repeat-containing protein (Kazama & Toriyama, 2014; Tang et al., 2014). The cloning of *Rf3* has not been reported.

In this study, we used an RT102-type CMS derived from wild rice (*Oryza rufipogon*) (Motomura et al., 2003). The mitochondrial genome of the RT102-type CMS (RT102A) harbors *orf352*, a sequence variant of *WA352* whose encoded protein differs from WA352 by four amino acid substitutions. The surrounding genomic structure at *orf352* in RT102A is also different from that of WA-type CMS. In WA-type CMS, the mitochondrial genome downstream of *WA352* contains exons 4 and 5 of *nad5*, encoding NADH dehydrogenase, whereas the corresponding sequence in RT102A originates from exons 1 and 2 of *rps3*, encoding ribosome small subunit 3. The shrunken pollen phenotype we observed in RT102-type CMS is similar to that of the WA-type CMS, although some spherical pollen grains are evident in RT102A (Okazaki et al., 2013). These signs of inhibition of pollen development are likely to be associated with microspore abortion at the early microspore stage, possibly via an identical mechanism in the RT102-type CMS and WA-type CMS. Genomic manipulation of *orf352* would validate its role as a CMS-causative gene, but the direct transformation of the rice mitochondrial genome has not been reported. As an alternative technique, we recently developed mitoTALENs; these are genome editors targeted to mitochondria, but expressed from a transgene in the nuclear genome. We used mitoTALENs to edit the mitochondrial gene *orf79* to demonstrate that disruption of *orf79* restores fertility of BT-type CMS rice (Kazama et al., 2019). Here, we tested the contribution of *orf352* to RT102-type CMS using two mitoTALEN constructs targeting *orf352*, culminating in the isolation of *orf352*-edited plants (Fig. 1B and C).

We previously showed that DSBs introduced by mitoTALENs around *orf79* in the mitochondrial genome are repaired via ectopic homologous recombination (Kazama et al., 2019). Next, using fusion primer and nested integrated PCR (FPNI-PCR), we determined that DSBs introduced by *orf352*-targeted mitoTALENs were repaired similarly. FPNI-PCR also allowed us to characterize the recombination sites in detail (Fig. 2 and Fig. S5). In all transformants, DSBs at the 5ʹ site involved the same homologous recombination event at the *orf352* sequence, which is identical to *orf284* and *orf284* itself. Repair of DSBs at the 3ʹ site was more variable and used various sequences of variable length, from 13 bp to 100 bp. Interestingly, the length of the 5ʹ recombination repair tract was the greatest, at 559 bp, and was common to all transformants. This observation suggests that the longer and highly similar stretches of sequences make for efficient templates for repair via homologous recombination. We classified all transformants into six types based on recombination at the 3ʹ site. Recombination resulted in complete loss of *orf352* in Type 1, 2, and 3 plants, but only partial loss, with retention of the portion of *orf352* that is homologous to *orf288*, in Type 4, 5, and 6 plants. Genome editing of the *orf352* partially rescued the pollen defects observed in RT102A, as evidenced by spherical pollen grain rich in starch seen in Type 1, Type 2, and Type 3 plants that can germinate on stigmas. Seed set, however, was not restored. We hypothesized that early microspore development is affected by *orf352*, while pollen and pollen tube growth may be influenced by new genes such as *orf284*, which are now duplicated in the mitochondrial genomes of our *orf352*-edited plants, or by other unidentified gene(s) in the RT102-type mitochondrial genome.

The partial restoration of pollen viability and development in Type 1 and 3 plants, as demonstrated by the accumulation of mostly spherical and darkly stained pollen grains with some germination potential, may be associated with the loss of *orf352*. Supporting this hypothesis, Type 5 and 6 plants showed no recovery of pollen development and retained a large segment of *orf352* sequence (encoding 179–352 aa of ORF352) homologous to *orf288* to yield the new mitochondrial gene *orf174*. This new gene may be expressed in the anther within spikelets at the meiotic stage. The mitochondrial genome of Type 4 plants harbored the new gene *orf142*, which encodes a protein comprising 211–352 aa of ORF352. Because these plants showed partially restoration of pollen development, unlike Type 5 and 6 plants, we propose that the 32 amino acids from ORF352, which are included in ORF174 but absent from ORF142 (Fig. 5), might be crucial for inhibition of pollen development, resulting in the production of shrunken and unstained pollen grains. A previous study of the function of WA352 in WA-type CMS reported that WA352 contains two COX11-interacting domains, from 218–292 aa and from 294–352 aa; in addition, interaction between COX11 and WA352 was required to establish CMS (Luo et al., 2013). The region spanning 169–199 aa, named the cs2 region, was also proposed to be a crucial in regulating the interaction between WA352 and COX11 (Tang et al., 2017). This region overlaps with the 32-aa segment we identified here (Fig. 5). The deletion of this region in ORF142 encoded by the mitochondrial genome of Type 4 plants might therefore prevent the interaction between ORF142 and COX11, leading to partial recovery of pollen development, although ORF142 is thought to still contain one COX11-interacting region. Further investigation of the function of these truncated ORF352 variants is required, in the context of their interaction with COX11. This report suggests that multiple genes in the mitochondrial genome may be involved in the induction of male sterility in rice. Our study also provides new insights into the function of CMS-causative genes and illustrate the usefulness of mitoTALENs for functional research of plant mitochondrial genes.

**Figure 5.**
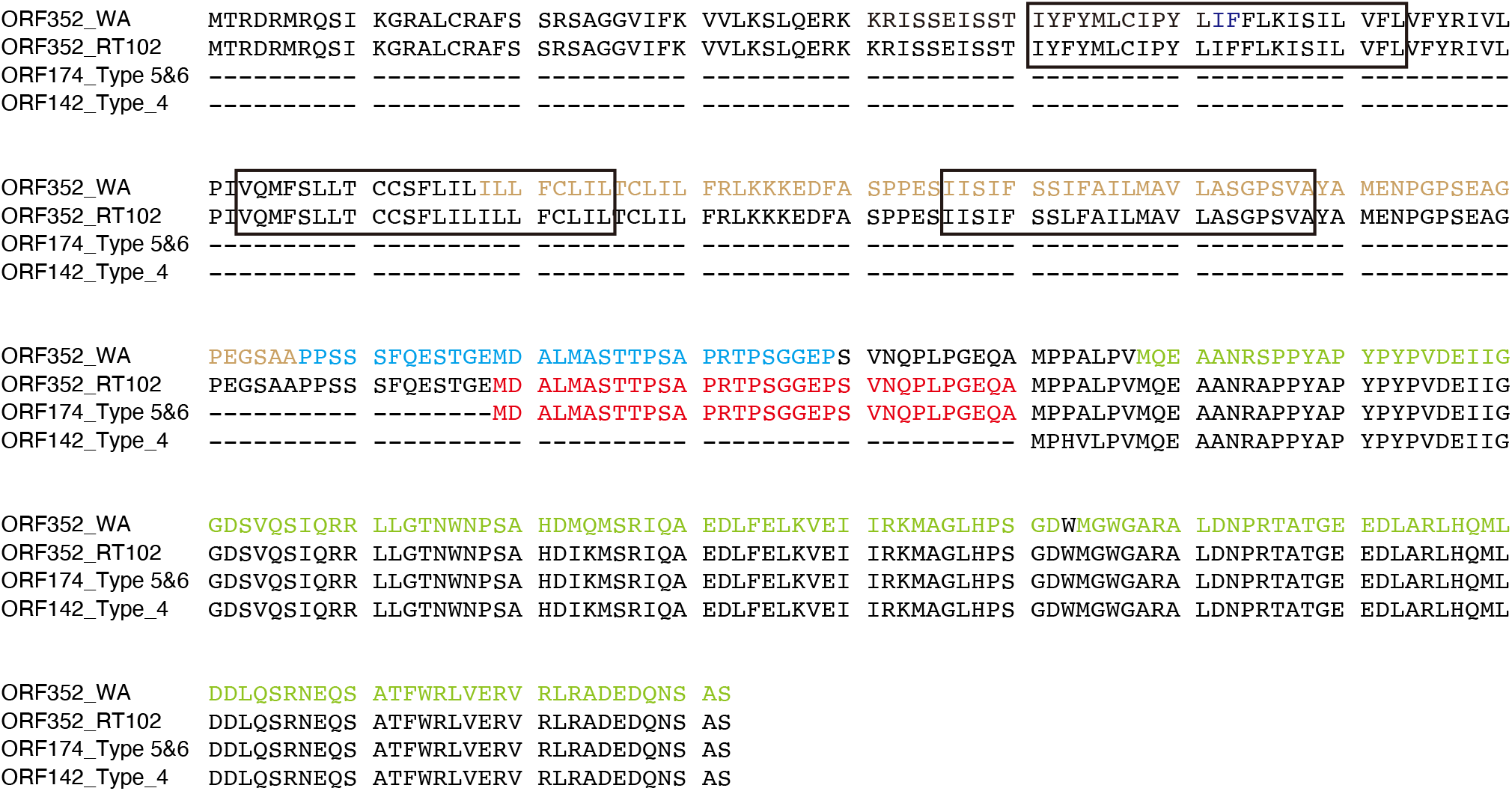
Alignment of ORF352, ORF174, and ORF142 protein sequences identified a 32-amino acid stretch that is critical to inhibition of pollen development. Brown, CS3 domain; blue, CS2 domain; green, COX11-interaction region (Tang et al., 2017); red, critical stretch of 32 amino acids for ORF352 function. Boxes indicate predicted transmembrane domains.

## Materials and methods

### Plant materials

The RT102-type CMS line (RT102A) was developed via multiple backcrosses between *Oryza rufipogon* Griff. accession W1125 (National Institute of Genetics, Mishima, Japan) and *Oryza sativa* L. cv. Taichung 65, as previously described (Okazaki et al., 2013). This RT102-type CMS comprises the nucleus of *Oryza sativa* L. cv. Taichung 65 and the cytoplasm of *Oryza rufipogon* Griff., accession W1125.

### Construction of mitoTALEN vectors

MitoTALEN binary vectors used here were constructed as described previously (Kazama et al., 2019). Briefly, DNA recognition motifs for the selected target sequences were assembled via the Golden Gate cloning, using methods with the Platinum Gate TALEN kit (Sakuma et al., 2013), into the multisite-entry vectors for the right or the left TALEN open reading frames (ORFs). ORFs in the entry vectors were then recombined into Gateway destination vectors harboring the cauliflower mosaic virus (CaMV) 35S promoter and the mitochondrial localization signal (MLS) from *Arabidopsis* ATPase deltaʹ subunit (encoded by At5g47030) by multi-LR reactions. The destination vector is based on the binary vector pH7WG (Karimi et al., 2002). An additional Gateway Entry vector, containing a transcription termination sequence from *Arabidopsis* heat-shock protein (HSP18.2; encoded by At5g59720), one copy of the 35S promoter followed by one copy of the MLS, was recombined into the destination vector between the right and left region of the TALEN ORFs to provide the transcript terminator for the TALEN right transgene, and the promoter and MLS for TALEN left (Fig. S1).

### Generation of transgenic plants with mitoTALEN vectors

Plants were planted in plastic pots filled with rice culture soil and grown at 30°C/25 °C (10-h day/14-h night) in a biotron (Nippon Medical & Chemical Instruments Co. Ltd, Osaka, Japan). Transformation methods were described previously (Kazama et al., 2008). Primary (T_0_) transformants were confirmed by PCR using a primer pair that amplifing a fragment of *Hygromycin Phosphotransferase* (*HPT*) (Table S1). *HPT*-positive T_0_ plants were transplanted to soil and transferred to a biotron. We obtained six transgenic plants with the TAL1 construct (1a, 1b, 2a, 5a, 6a, and 6b) and 12 with the TAL2 construct (1a, 1b, 2b, 3a, 3b, 3c, 4a, 4b, 5a, 5b, 6b, and 6c). The number of each plant reflects the petri plate from which the original T_0_ seedlings were selected. Plants with the same number but a different letter originated from the same petri plate. When T_0_ plants started to set seeds, heading panicles were bagged to avoid cross-pollination. At maturity, filled and unfilled spikelets were counted and seed setting rate was calculated. Spikelets were harvested 1 day before anthesis to stain pollen grains with a 1% (w/v) I_2_-KI Lugol solution to determine pollen viability. Pollen germination was observed on stigmas via aniline blue staining, as previously described (Fujii & Toriyama, 2005).

### Isolation of genomic DNA and Southern blot analysis

Total DNA was extracted from green leaf blades using the DNeasy Plant Mini Kit (Qiagen). The structure of the mitochondrial genome around the mitoTALEN target sequences was determined by PCR using the primers listed in Table S1, followed by sequencing to define the exact recombination site. Briefly, fusion primer nested integrated-PCR (FPNI-PCR) was performed with the primers listed in Table S1. The resulting PCR amplicons were purified after gel electrophoresis and sequenced. Recombination sites in the mitochondrial genome obtained by this method were confirmed by PCR and by Southern blot analysis, as described previously (Kazama et al., 2016).

## Acknowledgements

We thank M. Ito (Tohoku University) for technical assistance with TAL transformation. This research was partly supported by grants from the Japanese Science and Technology Agency (PRESTO to S. A.) and the Japan Society for the Promotion of Science (grant numbers JP16H06182, JP18H02172, and JP20H05680 to TK., and JP19H02927 to S.A.).

---

**Figure S1. Introduction and confirmation of the *orf352*-targeting mitoTALEN vectors, pTAL1 and pTAL2.**

**(A)** Schematic diagram of the T-DNA region from the mitoTALEN vectors. 35S pro, cauliflower mosaic virus 35S promoter; MLS, mitochondrial localization signal; TALEN Right, right side domain of TALEN; Ter, transcriptional terminator of *Arabidopsis* heat shock protein (HSP18.2; encoded by At5g59720), TALEN Left, left side domain of TALEN; HPT, *Hygromycin Phosphotransferases* gene. **(B)** Confirmation of stable transformation with pTAL1 and pTAL2 by genomic PCR. *Tubulin* serves as a positive control for amplification.

**Figure S2. Repair principle of the double-strand break introduced by mitoTALENs in the mitochondrial genome.**

A double-strand break introduced by a mitoTALEN will be repaired by homologous recombination. At the 5ʹ (or 3ʹ) side of the target site, the break is repaired between recombination site A (or B) near the target site and recombination site Aʹ (or Bʹ), which may be anywhere in the mitochondrial genome but is homologous to recombination site A (or B). As a consequence of repair by homologous recombination, the region between recombination sites A and B is deleted from the mitochondrial genome. Homologous recombination occurs non-reciprocally and is accompanied by replication, so finally, DNA molecules containing the site A/Aʹ and B/Bʹ will be duplicated.

**Figure S3. Detection of the deleted region around the *orf352* after homologous recombination by PCR.**

**(A)** Genomic structure around the *orf352* open reading frame. Black lines, from 1–9, indicate PCR amplicons. **(B)** PCR analysis of each transgenic plant for the nine genomic regions indicated in (A). T65 is a fertile *japonica* rice, which lacks the sequence around *orf352*. RT102A serves as a positive control for the presence of *orf352*. Primers are listed in Table S1.

**Figure S4. Schematic diagram of double-strand break repair in each type of transgenic plants.**

**(A)** Genomic organization of double-strand break repair at the 5ʹ side of the target site in all *orf352*-edited plants. Top, genomic structure around the *orf352* open reading frame. Pink, region identical to *orf284*; green, region homologous to *orf288*; orange, *orf352*-specific sequences. Scissors indicate mitoTALENs (TAL1 and TAL2). Middle, genomic structure around the recombination site Aʹ (illustrated in Fig. S2). Bottom, genomic structure of new recombinants. **(B–F)** Genomic organization of double-strand break repair at the 3’ side of the target site in Type 1 (**B**), Type 2 (**C**), Type 3 (**D**), Type 4 (**E**), and Type 5 or Type 6 (**F**) *orf352*-edited plants. Top, genomic structure around the *orf352* open reading frame. Pink, region identical to *orf284*; green, region homologous to *orf288*; orange, *orf352*-specific sequences. Scissors indicate mitoTALENs (TAL1 and TAL2). Red box indicates the recombination site B. Middle, genomic structure around the recombination site Bʹ (Fig. S2). Bottom, genomic structure of new recombinants. Black line indicates the region originating around *orf352*. Blue line indicates the region originating around the recombination site Bʹ. A new ORF is predicted at the recombination site for each recombinant type: o*rf77* (Type 1); *orf47* (Type 2); *orf108* (Type 3); *orf142* (Type 4); *orf174* (Types 5 and 6). Type 5 recombination also harbor sequences originating around an additional recombination site.

**Figure S5. Double-strand breaks induced by mitoTALENs are repaired by the identical homologous recombination at the 5ʹ side of the target site.**

**(A)** Top, genomic structure around the *orf352* open reading frame. Pink, region identical to *orf284*; green, region homologous to *orf288*; orange, *orf352*-specific sequences. Scissors indicate mitoTALENs (TAL1 and TAL2). Bottom, position (in bp) of recombination site A across all *orf352*-edited plants. Red boxes indicate the recombination sites A. **(B)** Sequence comparisons between a recombinant sequence and its corresponding sequence in the RT102-type mitochondrial genome.

